# Brazilin is a Natural Product Inhibitor of the NLRP3 Inflammasome

**DOI:** 10.1101/2023.10.30.564348

**Authors:** Emily McMahon, Sherihan El-Sayed, Jack Green, Christopher Hoyle, Lorna Fitzpatrick, Emma Jones, Eve Corrie, Rebecca L. Kelly, Mairi Challinor, Sally Freeman, Richard A. Bryce, Catherine B. Lawrence, David Brough, Paul R. Kasher

**Author notes:** **Corresponding author: Dr Paul Kasher, **.

## Abstract

Excessive or aberrant NLRP3 inflammasome activation has been implicated in the progression and initiation of many inflammatory conditions; however, currently no NLRP3 inflammasome inhibitors have been approved for therapeutic use in the clinic. Here we have identified that the natural product brazilin effectively inhibits both priming and activation of the NLRP3 inflammasome in cultured murine macrophages, a human iPSC microglial cell line and in a mouse model of acute peritoneal inflammation. Through computational modelling, we predict that brazilin can adopt a favourable binding pose within a site of the NLRP3 protein which is essential for its conformational activation. Our results not only encourage further evaluation of brazilin as a therapeutic agent for NLRP3-related inflammatory diseases, but also introduce this small-molecule as a promising scaffold structure for the development of derivative NLRP3 inhibitor compounds.

## Introduction

Inflammasomes are cytosolic, multimeric protein complexes which assemble within cells in response to danger signals evoked by tissue injury and/or infection. Inflammasomes serve a dual purpose of facilitating the maturation and release of pro-inflammatory cytokines interleukin (IL)-1β and IL-18 and initiating a lytic form of cell death known as pyroptosis. Several inflammasomes have been identified, each named after a unique pattern-recognition receptor (PRR) protein which recognises pathological stimuli and initiates inflammasome assembly. Such PRRs include absent-in-melanoma 2 (AIM2), pyrin and members of the nucleotide-binding oligomerization domain (NOD), leucine-rich repeat (LRR)-containing protein family (‘NLR’); NLRP1, NLRP3, NLRP6 and NLRC4 [1]. The NLRP3 inflammasome is the most widely characterised of all inflammasomes and has been the focus of our current research.

Canonical NLRP3 inflammasome activation is governed by a dual signalling mechanism. The first ‘priming’ signal involves extracellular toll-like receptor (TLR) or cytokine receptor activation, which initiates nuclear factor-κB (NF-κB)-dependent transcription of the NLRP3 protein and several pro-inflammatory cytokines including pro-IL-1β [2, 3]. A second ‘activation’ stimulus is then required to trigger inflammasome assembly. A range of damage-or pathogen-associated molecular patterns (DAMPs/PAMPs, respectively) have been found to activate the NLRP3 inflammasome, including crystalline/particulate substances (e.g. uric acid crystals or silica), extracellular ATP [2, 4] and signals associated with viral [5], fungal [6] and bacterial [7] infection. These activating stimuli evoke various cellular stress responses, including K^+^ efflux, calcium influx, reactive oxygen species production, mitochondrial DNA release, lysosomal destabilisation [8, 9] and a disruption of endosomal trafficking [10], which ultimately trigger the oligomerisation of NLRP3 proteins. This provides a platform for the polymerisation of apoptosis-associated speck-like protein containing a CARD (ASC), creating filament amalgamations known as ASC specks, which are considered a hallmark of inflammasome activation. Inflammasome assembly is complete with the recruitment of pro-caspase-1 enzymes, which undergo proximity-induced autocatalytic activation to form mature caspase-1 proteases [8, 9]. Caspase-1 then catalyses the maturation of IL-1β and IL-18 cytokines and the cleavage of gasdermin-D (GSDMD) proteins. GSDMD N-terminal fragments insert into the cell membrane and create pores, facilitating the extracellular release of mature cytokines [11, 12]. Finally, an oligomerisation of ninjurin-1 (NINJ1) cell-surface proteins is necessary to drive plasma membrane rupture and death of the cell [13–16].

Appropriate inflammasome activation and production of pro-inflammatory IL-1β is an essential part of a protective host immune response, however, dysregulation of such a response can potentiate or even initiate disease pathologies. Aberrant and/or excessive NLRP3 inflammasome activation has been implicated in several conditions, including diabetes, cryopyrin-associated periodic syndromes, neurodegenerative diseases such as Parkinson’s and Alzheimer’s disease, as well as both ischaemic and haemorrhagic stroke [17–19]. Inhibition of the NLRP3 inflammasome is a promising therapeutic strategy to attenuate inflammation-associated pathogenesis of such life altering and often under-treated conditions. MCC950, otherwise known as CRID3 or CP-456,773, is one of the first known and well-studied small-molecule inhibitors of the NLRP3 inflammasome [20, 21]. MCC950 potently and specifically inhibits NLRP3 inflammasome formation by binding to a site within the NLRP3 NACHT domain. This binding stabilises NLRP3 and prevents essential conformational changes required for NLRP3 activation [22–24]. However, concerns surrounding liver toxicity have prevented any clinical application of MCC950 [18]. As such, the development of new, safe NLRP3 inflammasome inhibitors remains an active and evolving area for drug discovery. There is now a growing library of NLRP3 inhibitor compounds which differ in their mechanisms of action and potential clinical applicability, with some currently in or having completed early-phase clinical trials [25].

Bioactive compounds from plants present an important avenue for drug discovery, with such natural products potentially carrying a lower risk of severe adverse effects and lower developmental costs relative to synthetic compounds. Brazilin (Figure 1A) is a small-molecule isoflavonoid (MW 286.28) which occurs naturally as the (S) stereoisomer (reference to brazilin here refers to this isomer). This compound yields a red coloured brazilein product upon oxidation (Figure 1B). Brazilin is highly abundant in the heartwood of several tree species, including brazilwood (*Caesalpinia echinate*), sappan (*Caesalpinia sappan*) and Mexican logwood (*Hematoxilum brasiletto*) [26]. Heartwood extracts from these trees have been widely used as colorants in foods, fabrics and cosmetics and have long been used as traditional remedies for a range of conditions, including dermatological problems, diabetes, fever and cancer [27, 28]. Preclinical research has described several pharmacological activities of brazilin, including an ability to attenuate the aggregation and cytotoxicity of amyloid beta protein [29], limit cancer cell proliferation [30] and reduce inflammation. Brazilin has been shown to reduce the production of pro-inflammatory mediators by cultured macrophages in response to bacterial lipopolysaccharide [31–33] and to exert an anti-inflammatory effect in rodent models of renal injury, significantly decreasing Tumour Necrosis Factor (TNF) α and IL-1β expression in the tissue [34] and increasing anti-inflammatory cytokine production [35].

**Figure 1.**
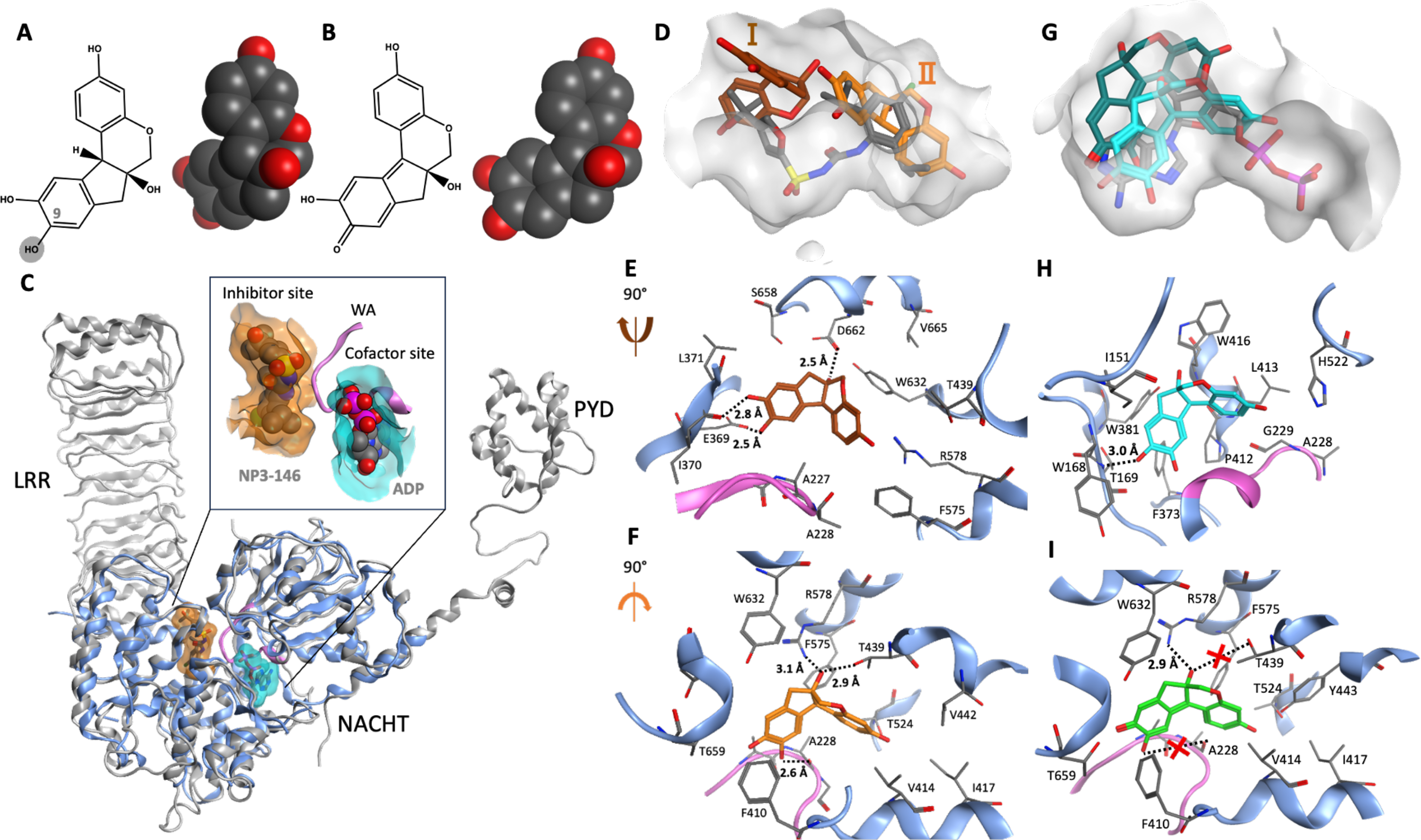
Predicted NLRP3 binding mode of brazilin from docking and simulation. 2D and 3D structures of (A) (S)-brazilin in which the grey highlighted OH group represents the hydroxyl group that oxidizes to give (B) (S)- brazilein. (C) Superposition of full-length NLRP3 cryo-EM structure (grey, PDB code 7PZC) [23] with X-ray structure of NACHT domain (blue, PDB code 7ALV) [36]; inset shows in more detail the two binding sites in NACHT domain. Walker A (WA) motif is coloured pink. (D) Superposition of X-ray pose of NP3-146 (grey), and top-ranked docked pose I (brown) and pose II (orange) of brazilin in inhibitor binding pocket (grey surface) of NLRP3-NACHT structure. Detailed interatomic interactions of (E) pose I (brown) and (F) pose II of brazilin (orange) with inhibitor binding site using NACHT-NLRP3 structure after 40 ns MD simulation. (G) Superposition of X-ray pose of ADP (grey), post-MD structure of brazilin (cyan) and brazilein (dark cyan) in cofactor binding pocket (grey surface) of NLPR3-NACHT structure. (H) Detailed interatomic interactions of brazilin (cyan) with cofactor binding site using NACHT-NLRP3 structure after 40 ns MD simulation. (I) Detailed interatomic interactions of pose II of brazilein (green) in the inhibitor binding site using the NACHT-NLRP3 structure after 40 ns MD simulation. Dashed lines represent hydrogen bonds and lost interactions indicated by dashed line with red cross. See also Video S1 showing the MD simulation of interactions between S-brazilin and the inhibitor binding site of NACHT domain.

However, to the best of our knowledge, the specific effect of brazilin on inflammasome responses has not yet been investigated.

Here we report that brazilin is an inhibitor of the NLRP3 inflammasome. Molecular modelling and simulation proposes that brazilin can adopt a favourable pose in the inhibitor binding site of the NLRP3 NACHT domain; the same site to which the selective NLRP3 inhibitor MCC950 binds to hold NLRP3 in an inactive conformational state. We show that brazilin can significantly inhibit the NLRP3 inflammasome response in primary murine bone marrow-derived macrophages (BMDMs) and induced pluripotent stem cell (iPSC)-derived human microglia. We also report that brazilin treatment can significantly attenuate the NLRP3 inflammasome response *in vivo*, using a mouse model of acute peritoneal inflammation, highlighting the potential effectiveness of using brazilin for the development of new therapeutics targeting NLRP3-dependent disease.

## Materials and Methods

### Molecular modelling and simulation

A complete NACHT domain structure of NLRP3 was prepared for computational docking from its X-ray structure (PDB code 7ALV) [36] using SWISS-MODEL server (https://swissmodel.expasy.org). Docking was performed using the OpenEye software suite [37]. The *Omega classic* tool was used to create 3D structures of brazilin and brazilein using a maximum of 500 conformations for each compound. The NLRP3 cofactor site and inhibitor site were prepared for docking using the *make_receptor* module. The *FRED* module was used to dock compounds using the Chemgauss4 scoring function. The best 50 poses for each compound were visualized using Vida 4.4.0 and MOE 2022.02 [38]. This docking protocol was previously validated and reported [39] by redocking of ADP in the cofactor binding site and NP3-146 (an analogue of MCC950) in the inhibitor site which reproduced the crystal structure pose well in both cases.

Nevertheless, molecular dynamics (MD) simulations [40–42] and experimentally determined structures [23, 24, 43–45] of NLRP3 point to the flexibility of the protein. Consequently, we refined our docked poses using MD simulation, following an approach recently applied by us in the analysis of tubulin inhibitors [46]. Therefore, MD simulations of NLRP3-ligand complexes were performed for selected docked poses. These calculations employed the AMBER 19 package [47]. Atomic partial charges of ligands were assigned *via* the AM1-BCC method using the *antechamber* module of AMBER. The *gaff2* [48] and *ff14SB* [49] force fields were used to describe ligands and receptor, respectively. Force field parameters for ADP were assigned from the AMBER database (http://amber.manchester.ac.uk/) [50] and *gaff2* [48]. The systems were solvated in an octahedral *TIP3P* water box [51] that extends at least 10 Å from the protein-ligand surface. The system was neutralized by addition of 2-5 chloride counterions to the solvated system. This led to ∼15,664 water molecules for each simulation system. The generated topology files were edited with the *parmed* [52] module of AMBER 19 to repartition the mass of heavy atoms into the bonded hydrogen atoms. The new topology file was designed to use hydrogen mass repartitioning [53] in which the time step of the simulation was 4 fs. The nonbonded cut-off of 9.0 Å was used, along with the particle mesh Ewald [54] method for long range electrostatic interactions.

MD simulations were performed using the *pmemd.cuda* module of AMBER 19. Before simulation, water molecules were relaxed by energy minimizing the solvated system. The system was heated to 300 K in two steps under NVT conditions (constant Number, Volume and Time) over 700 ps using the Langevin thermostat with a coupling constant value of γ = 2.0 [55]. Then the system was equilibrated at 300 K and 1 atm with a Monte Carlo barostat [56] for 10 ns with a coupling constant value of 25 and compressibility of 44.6. The production MD simulation was conducted for 40 ns in an NPT ensemble (constant Number, Pressure and Time), during which configurations were sampled every 10 ps.

For ligand-protein systems, the MM/GBSA method [57] was applied to the final 10 ns of the trajectory to estimate binding free energies. These calculations were performed using the *MMPBSA.py* [58] tool of AMBER 19. The internal and external dielectric constants were set to 1.0 and 80.0, respectively. The ionic strength was set to 0.15 mM. MM/GBSA calculations were performed using 100 snapshots/compound. The electrostatic contribution to binding free energy ΔG_el_ was a sum of electrostatic protein-ligand and solvation components; the non-electrostatic contribution ΔG_nonel_ was a sum of protein-ligand van der Waals and non-electrostatic solvation terms. Ligand efficiency (*LE*) was calculated by dividing the ΔG_tot_ by the molecular weight.

### Cell culture

#### Murine BMDMs

Primary BMDMs were harvested from C57BL/6J mice (aged 2-6 months, mixed sex) (Charles River). All procedures were carried out in accordance with the Home Office (Animals) Scientific Procedures Act (1986). Mice were euthanized by cervical dislocation. Bone marrow was isolated from the tibia and femur bones by cutting both ends of the bones and centrifuging them in an Eppendorf tube containing phosphate-buffered saline (PBS) at 10 000 g (10 s). The bone marrow was re-suspended in ACK lysis buffer (Lonza, LZ10-548E; Basel, Switzerland) to lyse the red blood cells. Remaining cells were passed through a strainer (70 µm pore size; Corning, 734-2761) before centrifugation at 1500 g (5 min). The cell pellet was resuspended in 70% Dulbecco’s Modified Eagle Medium (DMEM; Sigma-Aldrich D6429; St Louis, MO, USA); with the DMEM containing 10% (v/v) fetal bovine serum (FBS; Thermo, 10500064), 100 U ml^−1^ penicillin and 100 μg ml^−1^ streptomycin (PenStrep; Thermo, 15070063), supplemented with 30% L929 mouse fibroblast-conditioned medium. The BMDMs were cultured for 6-7 days in a humidified, 37 °C atmosphere with 5% CO_2_. Cells were then scraped and seeded overnight at a density of 1 × 10^6^ ml^−1^ in 24- or 96-well plates before being used for experiments.

#### Human iPSC-microglia

Human iPSC-microglia were differentiated according to protocols previously described using iPSC line KOLF2.1S obtained from the Wellcome Sanger Institute (HipSci) [59, 60]. Ethical approval details for the parental line can be obtained from https://hpscreg.eu/cell-line/WTSIi018-B-1. Briefly, iPSCs were grown in mTESR1 medium (STEMCELL Technologies, 85850; Vancouver, CA) on Geltrex dishes (Thermofisher, A1569601) until 80% confluent, then differentiated to macrophage precursors. iPSCs were seeded into 24-well Aggrewell 800 well plates (STEMCELL Technologies, 34811) to form embryoid bodies (EBS) in mTeSR1 supplemented with 50 ng ml^−1^ BMP-4 (Life Technologies, PHC9531), 50 ng ml^−1^ VEGF (Thermofisher, PHC9394) and 20 ng ml^−1^ SCF (Miltenyi Biotec, 130-093- 991; Bergisch Gladbach, Germany). At day 5 of culture, EBs were transferred to T175 flasks containing X-VIVO15 (Lonza, 02-053Q) supplemented with 100 ng ml^−1^ M-CSF (Peprotech, 300-25; London, UK), 25 ng/mL IL-3 (PHC0031), 2 mM Glutamax (35050061), and 50 μM β-mercaptoethanol (31350-010) (all Thermofisher) to generate macrophage precursors. Following 3 weeks of culture, macrophage precursors in the media supernatant were harvested weekly and differentiated into microglia-like cells using advanced DMEM/F12 supplemented with N2 (Thermofisher, 17502-048), 100 ng/mL IL-34 (Peprotech, 200-34), 10 ng/mL GM-CSF (Thermofisher, PHC2011), 2 mM Glutamax and 50 μM β-mercaptoethanol. Microglia-like cells were fed 2-3 times per week (media as per above) and experiments were all conducted between 11- and 14-days post-differentiation.

### Dose-response analyses of brazilin

Primary murine BMDMs were seeded in 96-well plates overnight before use (1×10^6^ cells/ml). Cells were primed with lipopolysaccharide (LPS; *E.coli* O26:B6) (1 μg ml^−1^) (Sigma, L2654) in DMEM (10% v/v FBS, 1% v/v penicillin–streptomycin, 1% v/v pyruvate) for 4 hours, before the media was replaced with ‘treatment media’ constituting DMSO (1% v/v) (Sigma, D2650) or brazilin (≥ 98% purity; HPLC) (Merck, SML2132; Rahway, NJ, USA) (30, 10, 3, 1, 0.3, 0.1, 0.03 or 0.01 μM) in serum-free DMEM (1% v/v penicillin–streptomycin, 1% v/v pyruvate). After 15 min treatment incubation, NLRP3 was activated by adding nigericin (10 µM) (Sigma, N7143) into the wells, with ethanol (0.5% v/v) used as a vehicle control, and incubated for 2 hours. Alternatively, to assess the direct effect of brazilin treatment on AIM2 or NLRC4 inflammasome activation, cells were transfected with either 1 μg ml^−1^ poly(deoxyadenylic-deoxythymidylic) acid sodium salt (poly(dA:dT), Sigma, P0883) or 1 μg ml^−1^ flagellin from *Salmonella typhimurium* (Invivogen, tlrl-stfla), respectively, using Lipofectamine 3000 (Thermofisher, L3000001) per manufacturer’s instructions, for 2 hours. Following 2-hour inflammasome activation, supernatants were collected and used for cell death analysis and Enzyme-Linked Immunosorbent Assays (ELISA).

### Inflammasome assays with brazilin

#### Primary BMDMs

To assess the effect of brazilin pre-treatment on LPS-induced NLRP3 inflammasome priming and nuclear factor erythroid 2-related factor 2 (NRF2) expression, primary BMDMs were pre-treated with brazilin (10 μM), the NRF2 inducer dimethyl fumarate (DMF) (30 μM) (Sigma, 242926) or DMSO vehicle (0.3 % v/v) in serum-free DMEM (1% v/v penicillin–streptomycin, 1% v/v pyruvate) for 15 min. LPS (1 μg ml^−1^) or PBS control was then added directly to the wells and incubated for 4 or 6 hours, as indicated in the results.

To assess the direct effect of brazilin treatment on canonical NLRP3 inflammasome activation, primary BMDMs were primed with 1 μg ml^−1^ LPS in DMEM (10% v/v FBS, 1% v/v penicillin– streptomycin, 1% v/v pyruvate) for 4 hours, followed by pre-treatment with brazilin (10 μM), the NLRP3 inflammasome inhibitor MCC950 (CP-456773 sodium salt; Sigma, P20280) (10 μM) or DMSO vehicle (0.3 %, v/v) in serum-free DMEM (1% v/v penicillin–streptomycin, 1% v/v pyruvate) for 15 min. To induce canonical NLRP3 activation, 10 µM nigericin or 5 mM adenosine triphosphate (ATP) (Sigma, A2383) was added directly to wells for 1 hour, 300 µg ml^−1^ silica (U.S. Silica, MIN-U-SIL 15) was added for 4 hours, 75 μM imiquimod (InvivoGen, R837) was added for 2 hours, or 1 mM Leu-Leu-O-methyl ester (LLOMe) (Sigma, L1002) was added to the wells for 1 hour. Alternatively, following 15 min pre-treatment of LPS-primed (1 μg ml^−1^; 4 h) primary BMDMs with brazilin (10 μM), MCC950 (10 μM) or DMSO vehicle (0.3 %, v/v) in serum-free DMEM (1% v/v penicillin–streptomycin, 1% v/v pyruvate), media was changed for K^+^-free buffer (145 mM NaCl and 10 mM HEPES, pH 7.47) containing brazilin (10 μM), MCC950 (10 μM) or DMSO vehicle (0.3 % v/v) for 4 hours to induce NLRP3 activation directly via K^+^ efflux. A K^+^-containing buffer (145 mM NaCl, 5 mM KCl and 10 mM HEPES, pH 7.44) was used as control.

To assess whether the effect of brazilin on NLRP3 inflammasome activation was reversible, LPS-primed (1 μg ml^−1^; 4 h) primary BMDMs were treated with brazilin (10 μM), MCC950 (10 μM) or DMSO vehicle (0.3 %, v/v) in serum-free DMEM (1% v/v penicillin–streptomycin, 1% v/v pyruvate) for 15 minutes. Subsequently, treatment media was either left on the cells (15 min), removed and replaced with fresh DMEM without drug treatments (x2, 15 min), or removed and replaced with fresh DMEM (x1) before a final removal and replacement with fresh DMEM containing vehicle (DMSO), MCC950 (10 µM) or brazilin (10 µM) (15min). Nigericin (10 µM, 2 h) was then added directly to the wells to activate NLRP3.

At the end of these experiments, supernatants were collected and processed using cell death and/or ELISA analyses, while lysis buffer (50 mM Tris-HCl, 150 mM NaCl, Triton 1%; pH 7.5) was added to the cells. Alternatively, cells were lysed in-well by adding protease inhibitor cocktail (Merck Millipore, 539131) (1% v/v) and Triton-X-100 (1% v/v) (Sigma, X-100) directly into the culture medium, to assess total protein content in combined cell lysate and supernatant via Western blot (as indicated in the results). All samples were stored at -20°C between use.

#### Human iPSC-microglia

To examine inflammasome activation, iPSC-microglial cells were transduced with a lentivirus expressing hASC (NM-013258.5) fused to GFP at 8-days-*in vitro* (DIV). After 72 hours, cells were treated with or without LPS (100 ng ml^−1^) for 16 hours. Microglial cells were then treated with or without MCC950 (0.1 µM), brazilin (10, 30, 60 or 100 µM) or DMSO control as indicated for 30 minutes prior to stimulation with nigericin (10 µM) for 2 hours. Images were captured using SX5 incucyte imaging system (20X phase and green fluorescence) and speck analysis was performed by blinded manual counting of specks in two images for each biological replicate in each treatment condition. Media supernatant was collected from the cells at the end of the experiment and IL-1β protein was analysed using an IL-1β Human Luminex® Discovery Assay kit (R&D Systems, LXSAHM; Abingdon, UK).

### Assessing the effect of brazilin treatment on inflammasome activation *in vivo*

#### Animals and housing conditions

Adult male C57BL/6J mice (n=20) aged 8-12 weeks were raised in-house at the University of Manchester. Mice were group housed and maintained on a regular 12-hour light-dark cycle (07:00 to 19:00 light phase) with standard housing conditions (constant temperature (21°C) and humidity, *ad libitum* food and water and environmental enrichment). All experiments were performed in accordance with the UK Home Office regulations (PPL: P4035628) and reported according to the ARRIVE guidelines for experiments involving animals [61].

#### Treatment protocol and sample collection

A single dose of LPS (1 µg in 500 µL sterile PBS) was administered via intraperitoneal (i.p.) injection, immediately followed by a single i.p. dose of either brazilin (50 mg/kg in PBS, 1% (v/v) DMSO) (n=5), MCC950 (20 mg/kg in PBS) (n=5) or vehicle (PBS with 1% (v/v) DMSO) (n=5). The experimenter was blinded to drug treatments. After 2 hours the mice were stably anaesthetised (3% isoflurane in 64% N_2_O and 33% O_2_), before i.p. administration of ATP (100 mM in PBS (500 µL/mouse)) or PBS vehicle. After 15 minutes, peritoneal lavage was performed by injecting 3 ml sterile RPMI-1640 media (Sigma, R0883). Peritoneal lavage samples were stored at -80°C until analysed by ELISA for IL-1β, TNFα and IL-6 content. The experimenter was un-blinded to the drug treatments only after conducting statistical analyses.

### Cell death and ELISA analyses

Cell death was quantified using a lactate dehydrogenase (LDH) assay kit (Promega; Madison, WI, USA), according to the manufacturer’s instructions. IL-1β (DY401), IL-6 (DY406) and TNFα (DY410) were analysed by ELISA, according to the manufacturer’s instructions (DuoSet, R&D Systems). Absorbance readings were measured using a Synergy™HT plate reader (BioTek, Winooski, VT, USA).

### Western blot

Cell lysates were diluted in Laemmli buffer (5×) (20% v/v) and heated (95°C, 10 min) before loading into the gels. Alternatively, as indicated in the results, proteins were concentrated in the samples by mixing in-well lysates with an equal volume of trichloroacetic acid (Fisher, 10391351) and centrifuged at 4°C, 18,000 x g (10 min). The supernatant was discarded, and the pellet resuspended in 100% acetone before centrifuging at 4°C, 18,000 x g (10 min). The supernatant was removed, and the pellet left to air-dry before resuspending in Laemmli buffer (2×). Samples with equal amounts of protein were loaded into the gels.

Samples were assessed for NLRP3, pro-IL-1β, mature IL-1β, pro-caspase-1, caspase-1 p10, gasdermin D and NRF2. Samples were run on SDS polyacrylamide gels and transferred at 25 V onto PVDF membranes using a semi-dry Trans-Blot® Turbo Transfer™ System (Bio-Rad). Membranes were blocked with 5% BSA in phosphate-buffered saline, 0.1% Tween 20 (Sigma) (PBST) for 1 hour at room temperature. Membranes were then washed with PBST (3x, 15 min) before incubation with either mouse anti-mouse NLRP3 monoclonal antibody (1 µg ml^−1^; Cryo2, Adipogen, G-20B-0014-C100), goat anti-mouse IL-1β polyclonal antibody (250 ng ml^−1^; R&D Systems, AF-401-NA) rabbit anti-mouse caspase-1 + p10 + p12 monoclonal antibody (1.87 µg ml^−1^; Abcam, ab179515), rabbit anti-mouse gasdermin D antibody (0.6 µg ml^−1^; Abcam, ab209845) or rabbit anti-mouse NRF2 (1.5 µg ml^−1^; CST, 12721) in 0.1% (IL-1β, NLRP3, caspase, gasdermin D) or 2.5% (NRF2) BSA in PBST, overnight at 4°C. Membranes were again washed with PBST (3x, 15 min) before incubation with either rabbit anti-mouse IgG (1.3 µg ml^−1^, 1% BSA in PBST; Agilent, P026002-2), goat anti-rabbit IgG (250 ng ml^−1^, 1% BSA in PBST; Agilent, P044801-2) or rabbit anti-goat IgG (500 ng ml^−1^, 1% BSA in PBST; Agilent, P044901-2), for 1 hour at room temperature. β-Actin (Sigma, A3854) was the loading control, using a monoclonal anti-β-Actin-peroxidase antibody (Sigma). After washing with PBST (3x, 15 min), proteins were visualized by applying Amersham ECL Western Blotting Detection Reagent (GE Healthcare, RPN2236) to the membranes and imaging with the G:BOX Chemi XX6 (Syngene) and Genesys software. Densitometry was performed using Fiji (ImageJ).

### Statistical analyses

Statistical analyses were conducted using GraphPad Prism 10.0.3 (GraphPad Software Inc., San Diego, CA). All data are presented as mean ± standard error of the mean (S.E.M.). Statistical significance was accepted as p<0.05.

The dose-response curve showing IL-1β release in response to NLRP3 activation was fitted using a 4- parameter logistical sigmoidal model to obtain an IL-1β half-maximal inhibitory concentration (IC_50_) value for brazilin. All other data were assessed for normality using a Shapiro-Wilk test before performing further analyses. Normally distributed data were analysed using either a one-way ANOVA with Tukey’s post hoc analyses or two-way ANOVA with Tukey’s or Šídák’s post hoc analyses, as indicated in the results, to compare mean values between three or more groups. Alternatively, paired t-tests were used to compare mean values between two groups. Data not normally distributed were analysed using a Kruskal-Wallis test with Dunn’s post hoc analyses, to compare mean values between three or more groups. Data showing fold changes compared to controls were analysed using one-sample t-tests, comparing means to a hypothetical mean of 1.0. (representing the control group mean); ANOVAs could not be used as the data for control groups had no distribution.

## Results

### Molecular modelling predicts a direct interaction of brazilin with NLRP3

Firstly, we evaluated the potential for brazilin to bind favourably to the NLRP3 protein using computational modelling. Interestingly, due to its fused ring structure, the conformer of brazilin forms a rigid, twisted L-shaped structure (Figure 1A). For comparison, the binding modes of the oxidized form, brazilein (Figure 1B), were similarly characterized. Recent cryo-EM and X-ray structures of NLRP3 indicate the presence of a ligand binding site, adjacent to but distinct from the cofactor site, with which the inhibitor MCC950 or its analog NP3-146 interacts (Figure 1C) [22–24, 43]. To evaluate its potential to bind to NLRP3, brazilin was docked in both the cofactor and the inhibitor binding sites, with subsequent refinement by molecular dynamics simulation.

Brazilin was predicted via docking to bind to the inhibitor binding site in the NACHT domain of NLRP3 in two distinct binding modes, denoted I and II, occupying opposite ends of the site (Figure 1D). After MD simulation of the docked pose I, with ADP bound to the cofactor site, the OH groups of brazilin form hydrogen bonds with residues Asp662, Ile370 and Glu369 of NLRP3 (Figure 1E). MD simulation of pose II of brazilin optimized hydrogen bonds with Thr439 as well as Ala228 and Arg578; the latter two amino acids are identified as key amino acids for binding of MCC950 and its analogs in the NACHT domain X-ray structure of NLRP3 (Figure 1F and Video S1). These poses are predicted to be similar in affinity, based on computed binding free energies using these trajectories, with values of - 31.6 and -31.7 kcal/mol for poses I and II, respectively (Table 1). These values are rather lower in affinity than the predicted free energy of nanomolar ligand NP3-146 (Table 1); however, the similarity in ligand efficiency of NP3-146 and brazilin (Table 1) does suggest a favourable interaction of brazilin with the inhibitor site of NLRP3.

**Table 1.**
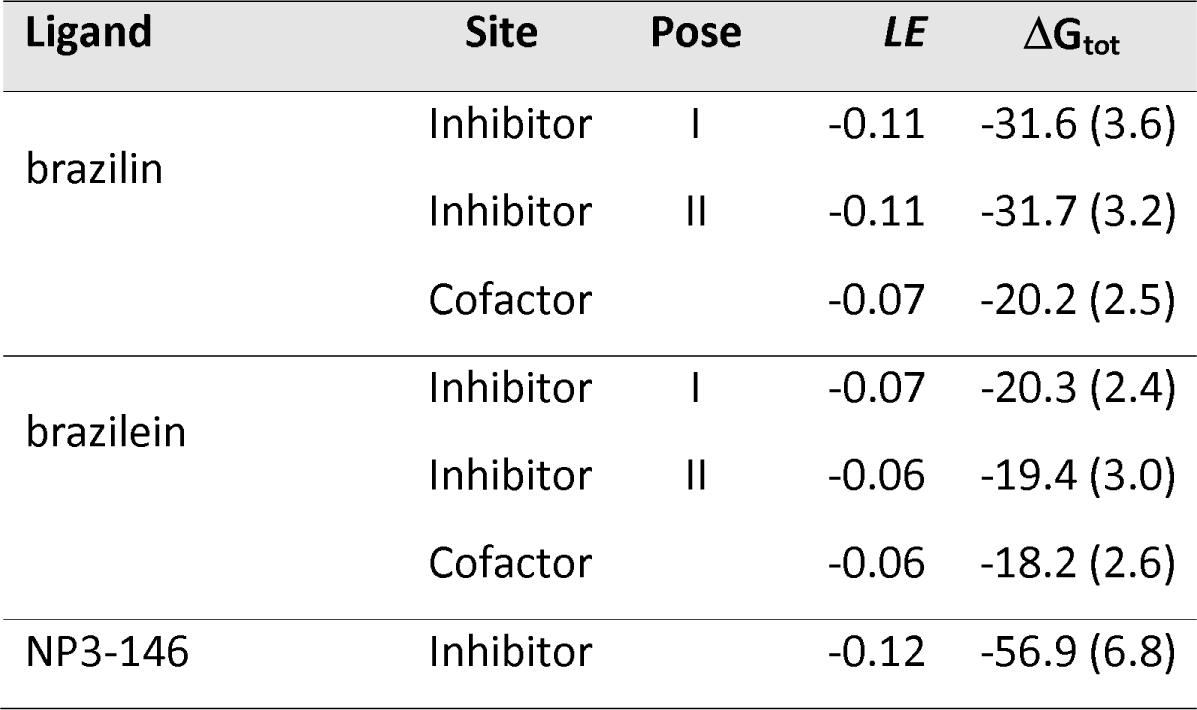
Computed binding free energies of brazilin and brazilein in the inhibitor and cofactor binding sites of the NLRP3 protein. Calculated total binding free energies (ΔG_tot_) are shown, using the MM/GBSA method for docked poses of brazilin and brazilein in inhibitor and cofactor binding sites of NLRP3. Binding free energy of X-ray pose of NP3-146 to NLRP3 inhibitor site also calculated. Energies in kcal/mol. Standard deviations in parentheses. Ligand Efficiency (*LE*) is ΔG_tot_/MW.

For comparison, brazilin was modelled into the cofactor binding site using the same docking/MD protocol. The ligand made a range of interactions in the site, including a hydrogen bond between an OH group of brazilin and the backbone N atom of Thr169 (Figures 1G,H); and several hydrophobic residue contacts (Table S1). However, the computed binding free energy for brazilin in this site was rather less than for the inhibitor site, with a value of -20.2 kcal/mol (Table 1).

Finally, the oxidised form of brazilin, namely brazilein, was docked into the two binding sites on the NLRP3 NACHT domain. The two MD-refined poses of brazilein in the inhibitor site exhibited a significant decrease in binding affinity relative to brazilin (Table 1). For pose I, flipping of brazilein occurred during the simulation and no hydrogen bonding was found (Table 1). For pose II of brazilein, a hydrogen bond formed between its OH group and Arg578 during molecular dynamic simulation (Figure 1I), but the additional hydrogen bonds formed with Ala228 and Thr439 by brazilin were lacking (Figure 1F). As for brazilin, the predicted binding energy of brazilein in the cofactor site was rather lower than that its interaction with the inhibitor site (Table 1), and 2 kcal/mol lower than for brazilin in the cofactor site (Table 1 and Figure 1G).

Therefore, our computational modelling suggests that the twisted L-shape of brazilin (Figure 1A) is capable of interacting well with the NACHT inhibitor site of NLRP3, in terms of shape complementarity and forming a hydrogen bonding network; however, the more planar curved shape of brazilein (Figure 1B) is predicted to be less compatible with either the cofactor or inhibitor sites. Therefore, we hypothesised that brazilin was a strong candidate as an inhibitor of the NLRP3 inflammasome.

### Brazilin inhibits activation of the NLRP3 but not AIM2 or NLRC4 inflammasomes in murine BMDMs

We next evaluated the ability of brazilin to inhibit the activity of NLRP3 experimentally using primary murine macrophages. LPS-primed (1 µg ml^−1^, 4 h) BMDMs were treated with brazilin (0.01-30 µM, 15 min) prior to the addition of nigericin (10 µM, 2 h) to activate NLRP3. Brazilin caused a dose-dependent inhibition of nigericin-induced IL-1β release, with an IC_50_ of 1.98 μM (Figure 2A_(i)_) and cell death also decreased in a dose-dependent manner (Figure 2A_(ii)_). Brazilin itself was determined as non-toxic to the cells, as treatment of either naïve cells or LPS-primed cells with brazilin in the absence of nigericin stimulation did not induce cell death at any concentration (Figure 2A_(ii)_).

**Figure 2.**
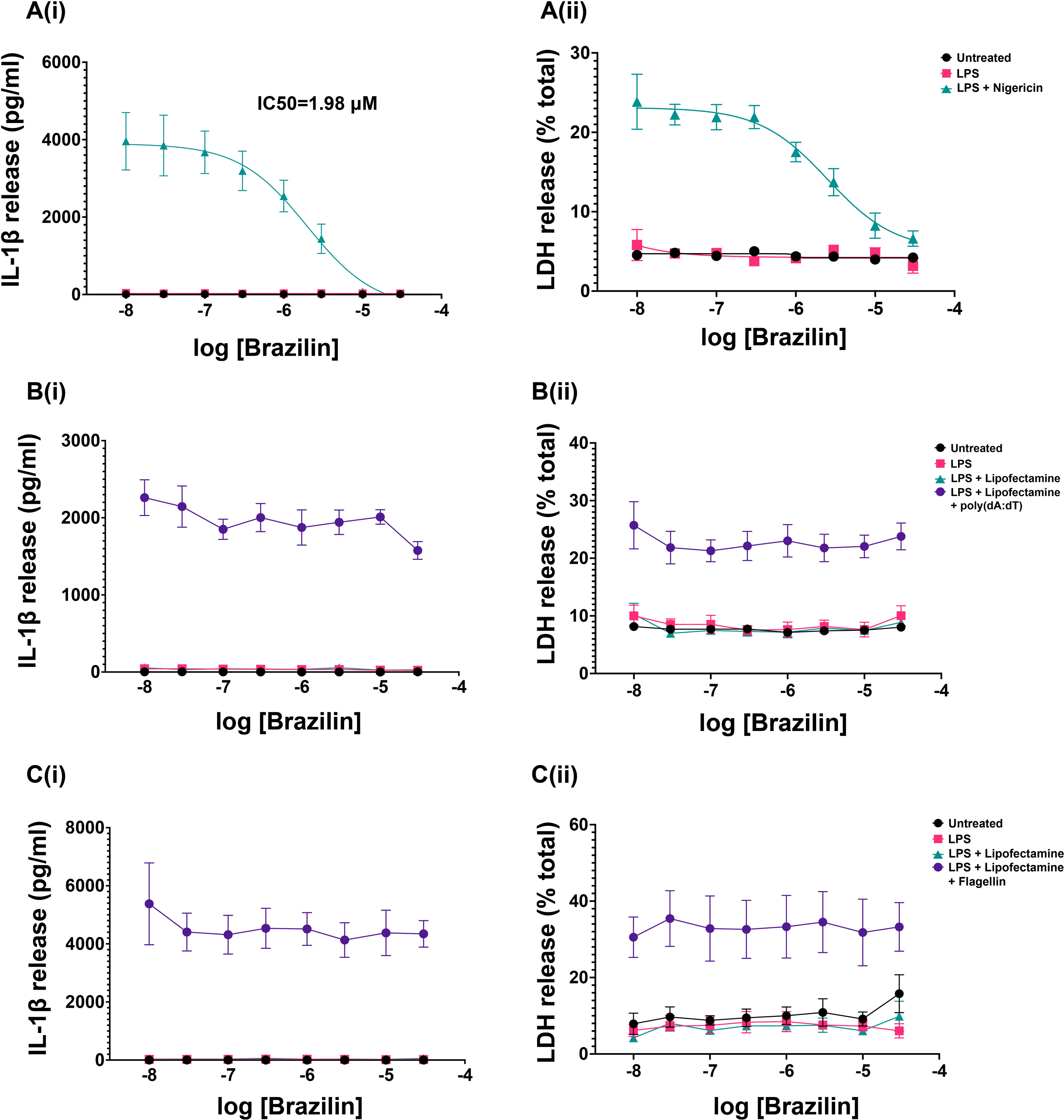
Brazilin causes a dose-dependent inhibition of the NLRP3 inflammasome but not AIM2 or NLRC4 inflammasomes, in primary murine BMDMs. LPS-primed (1 μg ml^−1^, 4 h) primary murine BMDMs were treated with brazilin at a range of concentrations (30, 10, 3, 1, 0.3, 0.1, 0.03, 0.01 μM) for 15 minutes, before (A) adding nigericin (10 µM, 2 h) to activate the NLRP3 inflammasome, (B) poly(dA:dT) transfection (1 μg ml^−1^, 2 h) to activate the AIM2 inflammasome, or (C) flagellin transfection (1 μg ml^−1^, 2 h) to activate the NLRC4 inflammasome. Supernatants were assessed for (i) IL-1β release by ELISA and (ii) lactate dehydrogenase (LDH) (cell death). To obtain an IL-1β half-maximal inhibitory concentration (IC_50_) value for brazilin against the NLRP3 inflammasome, the dose–response curve (Ai) was fitted using a 4-parameter logistical sigmoidal model. All data points are presented as mean ± SEM (N=3 biological repeats).

To deduce whether the inhibitory activity of brazilin was specific to the NLRP3 inflammasome, we also tested the effects of brazilin on NLRC4 and AIM2 inflammasome activation. Treatment of LPS-primed (1 µg ml^−1^, 4 h) BMDMs with brazilin (0.01-30 µM, 15 min) did not alter IL-1β release or cell death induced by subsequent transfection with poly(dA:dT) (AIM2 activator) (Figure 2B) or flagellin (NLRC4 activator) (Figure 2C). Collectively, these data indicate that brazilin can inhibit the activation of the NLRP3 inflammasome, but not AIM2 or NLRC4 inflammasome activation.

### Brazilin inhibits the activation step of the canonical NLRP3 inflammasome

As discussed previously, the NLRP3 inflammasome can be activated by a range of DAMPs and PAMPs. Despite having different upstream signalling pathways, most known stimuli ultimately cause K^+^ efflux as a necessary signal for NLRP3 activation. This includes nigericin (K^+^/H^+^ ionophore), ATP (P2X purinoceptor 7 agonist), silica (particulate material) and LLOMe (lysosome disrupting agent) [9]. However, the NLRP3 inflammasome can also be activated independently of K^+^ efflux using the imidazoquinoline compound imquimod [62]. We sought to investigate whether brazilin could directly inhibit NLRP3 inflammasome activation induced by these different stimuli, to address whether the inhibitory effect was dependent upon an interference with K^+^ efflux and/or was confined to disrupting a particular activation pathway.

We first confirmed a significant inhibition of nigericin-induced NLRP3 inflammasome activation by brazilin (10 µM) (Figure 3A_(i-ii)_), before testing brazilin at this concentration against other NLRP3 activating stimuli. Brazilin was also found to significantly inhibit other K^+^ efflux-dependent pathways of activation, namely activation by ATP (Figure 3A_(i-ii)_), silica (Figure 3B_(i-ii)_) and LLOMe (Figure 3C_(i-ii)_). K^+^ free buffer was also used to directly evoke cellular K^+^ efflux and NLRP3 inflammasome activation, in the absence of additional upstream signals, and under these conditions brazilin also significantly inhibited inflammasome activation (Figure 3D). Brazilin also inhibited K^+^ efflux-independent NLRP3 inflammasome activation in response to imiquimod (Figure 3E_(i-ii)_). Brazilin treatment in between LPS priming and nigericin or ATP stimulation prevented the cleavage of IL-1β and caspase-1 into their mature forms, and the cleavage of gasdermin D to produce its active N-terminal fragment (Figure 3F). Brazilin treatment did not affect pro-IL-1β expression but caused a slight reduction in NLRP3 levels when added after LPS priming.

**Figure 3.**
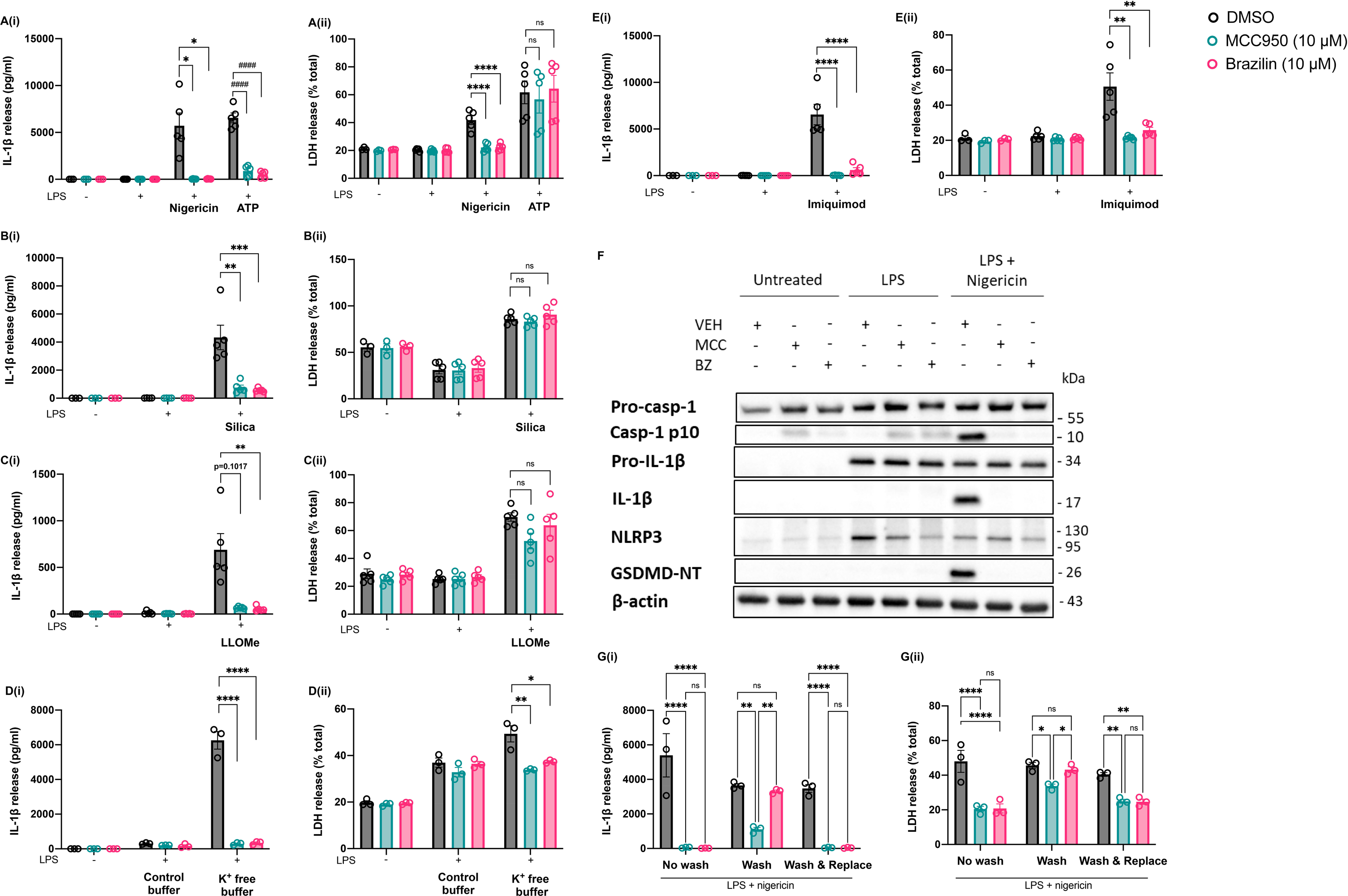
Brazilin inhibits both K efflux-dependent and -independent pathways of canonical NLRP3 inflammasome activation in murine macrophages. LPS-primed (1 µg ml^−1^, 4 h) primary murine BMDMs were treated with vehicle (DMSO), MCC950 (10 µM) or brazilin (10 µM) for 15 minutes. To activate NLRP3: **(A)** Nigericin (10 µM, 2 h) (N=5), ATP (5 mM, 2 h) (N=5), **(B)** silica (300 µg ml^−1^, 4 h) (N=5), **(C)** Leu-Leu-O-methyl ester (LLOMe; 1 mM, 1 h) (N=5), or **(E)** imiquimod (75 µM, 2 h) (N=5) was then spiked. Alternatively, **(D)** media was changed for K -free buffer containing DMSO, MCC950 (10 µM) or brazilin (10 µM) to activate NLRP3 (N=3). Supernatants were assessed for (i) IL-1β release by ELISA and (ii) lactate dehydrogenase (LDH) (cell death). **(F)** LPS-primed BMDMs were treated with DMSO, MCC950 or brazilin before nigericin stimulation, as detailed above. Concentrated protein content of the combined supernatant and cell lysates were probed for several markers of inflammasome activation by Western blotting. The blot shown is representative of 3 biological repeats. **(G)** Primed BMDMs (LPS 1 µg ml^−1^, 4 h) were treated with vehicle (DMSO), MCC950 (10 µM) or brazilin (10 µM) for 15 minutes. Over a further 15 minutes, cells either remained unaltered (‘no wash’), their media was removed and replaced with fresh media (x2) (‘wash’), or their media was removed (x2) and replaced with fresh media containing vehicle (DMSO), MCC950 (10 µM) or brazilin (10 µM) (‘wash & replace’). Nigericin (10 µM, 2 h) was then spiked to activate NLRP3 (N=3). Supernatants were assessed for (i) IL-1β release by ELISA and (ii) lactate dehydrogenase (LDH) (cell death). All data are presented as mean ± SEM, each data point (‘N’) representing a biological repeat. Statistical analyses following normality testing: (A-B^i^) ATP, silica, (D-E^i^) K free buffer, imiquimod, (A-E_ii_) Nigericin, silica, LLOMe, imiquimod and K free buffer data, (G): all one-way ANOVAs with Tukey’s post hoc comparisons. (A_i_) Nigericin, (C_i_) LLOMe, (A_ii_) ATP data: all Kruskal-Wallis tests with Dunn’s post hoc comparisons. *p < 0.05, **p < 0.01, **** or p < 0.0001. BMDMs, bone marrow-derived macrophages; DMSO, dimethylsulfoxide; LPS, lipopolysaccharide; VEH, vehicle (DMSO); MCC, MCC950; BZ, brazilin; Casp-1, caspase-1; GSDMD-NT, Gasdermin D N-terminal domain.

We next tested whether the inhibition of NLRP3 caused by brazilin was reversible. When brazilin was removed after 15 minutes treatment of LPS-primed (1 µg ml^−1^, 4 h) BMDMs, and not replaced before the addition of nigericin, IL-1β release and pyroptosis were no longer inhibited. However, when the cells remained exposed to brazilin for the duration of stimulation with nigericin (1 h), either when brazilin was not removed or brazilin was removed after 15 minutes treatment and then replaced before the addition of nigericin, inflammasome inhibition was maintained (Figure 3G_(i-ii)_). These data indicated that the inhibitory effect of brazilin on NLRP3 inflammasome activation was reversible.

Collectively, these results confirm an ability of brazilin to directly inhibit the activation step of the canonical NLRP3 inflammasome pathway, whether activation is dependent upon or independent of K^+^ efflux, and that the mechanism of inhibition is reversible.

### Brazilin can also inhibit the priming step of canonical NLRP3 inflammasome activation

As brazilin was effective at inhibiting the activation step of the NLRP3 inflammasome, we then investigated the role of brazilin on the priming step. To assess the potential effect of brazilin on the priming stage of the canonical NLRP3 inflammasome pathway, primary murine BMDMs were treated for 15 minutes with either DMSO (vehicle) or brazilin (10 µM) prior to the addition of LPS (1 µg ml^−1^, 4 h). Brazilin pre-treatment significantly reduced expression of pro-IL-1β, NLRP3 (Figure 4A) and IL-6 (Figure 4B_i_) in response to LPS priming. These data suggest that brazilin was also inhibiting priming of the canonical NLRP3 inflammasome. However, brazilin-mediated inhibition of priming was found to occur independently of its influence on the activation step of the pathway, as pro-IL-1β levels were only reduced when brazilin was added prior to but not after LPS priming (Figure 3F). In contrast, brazilin pre-treatment significantly enhanced LPS-induced TNFα release from the cells (Figure 4B_ii_). Cell death was not induced by either LPS stimulation or drug treatment (Figure 4B_iii_).

**Figure 4.**
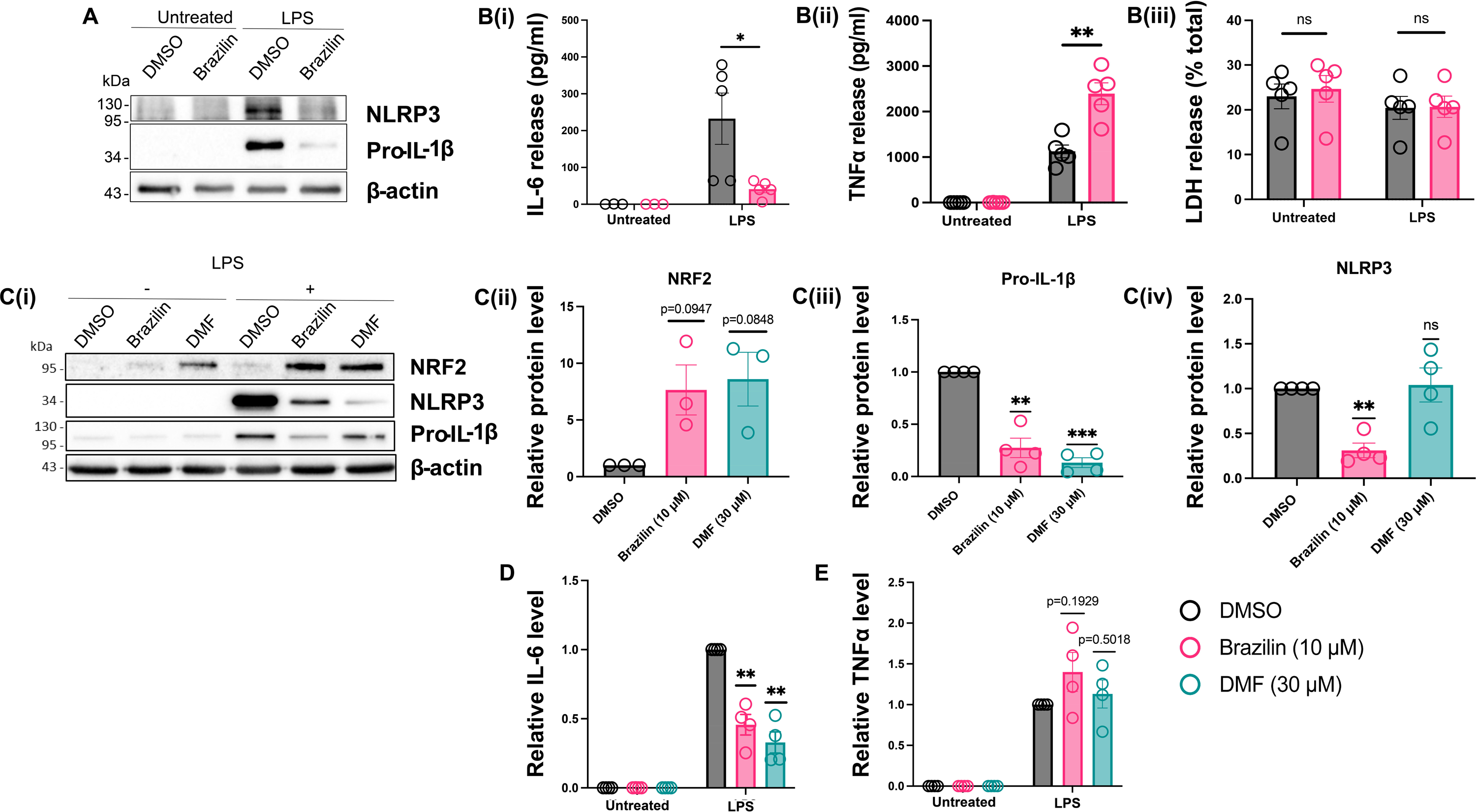
Pre-treatment with brazilin reduces LPS-induced priming of the canonical NLRP3 inflammasome in association with enhanced NRF2 expression. **(A-B)** BMDMs were treated with vehicle (DMSO) or brazilin (10 µM) for 15 min. LPS (1 µg ml^−1^, 4 h) was then added to the wells to induce priming. (A) Concentrated protein content from combined supernatant and cell lysates were probed by Western blotting for pro-IL-1β and NLRP3 proteins. The blot shown is representative of 3 biological repeats. **(B)** In a separate experiment, supernatants alone were analysed by ELISA for (B_i_) IL-6 (N=3 or 5), (B_ii_) TNFα (N=5) and (B_iii_) LDH release (N=5). **(C-E)** BMDMs were treated with vehicle (DMSO), brazilin (10 µM) or DMF (30 µM) for 15 min. LPS (1 µg ml^−1^, 6 h) was then added to the wells to induce priming. (C) Cell lysates were probed by Western blotting for NRF2, pro-IL-1β and NLRP3 proteins. (C_i_) The blot shown is representative of 4 biological repeats. For LPS+ samples only, expressions of (C_ii_) NRF2 (N=3), C(_iii_) pro-IL-1β (N=4) and C(_iv_) NLRP3 (N=4) proteins were quantified by densitometry (expressed relative to DMSO treatment). **(D-E)** Supernatants from the same cells were analysed by ELISA for (D) IL-6 (N=4) and (E) TNFα (N=4) release. Data show cytokine levels relative to DMSO + LPS treatment. See also Figure S1 for cytokine release data without normalisation. All data are presented here as mean ± SEM, each data point (‘N’) representing a biological repeat. Statistical analyses following normality testing: B(_i-iii_) paired t-Tests; C(ii-iv) and (D-E) one-sample t-Tests, comparing means to a hypothetical mean of 1.0. *p<0.05, **p < 0.01, ***p < 0.001. BMDMs, bone marrow-derived macrophages; DMSO, dimethylsulfoxide; DMF, dimethyl fumarate; NRF2; nuclear factor erythroid 2-related factor 2.

These data suggested that brazilin was affecting NF-κB response genes in a manner comparable to the effects of enhanced Nuclear factor erythroid 2-related factor 2 (NRF2) signalling. NRF2 is a transcription factor which is inactive when bound to Kelch-like ECH-associated protein 1 (KEAP1) in the cytoplasm. Some electrophilic agents and reactive oxygen species can induce KEAP1 degradation, allowing NRF2 to translocate into the nucleus where it can induce the expression of several target genes, but also supress the transcription of NF-κB secondary response genes, including IL-6 and IL-1β [63, 64]. NRF2 accumulation has a much lesser effect on NF-κB primary response genes, such as TNFα [64, 65]. Since brazilin supressed IL-6 and IL-1β production but not TNFα, we investigated whether brazilin could induce NRF2 signalling. BMDMs were pre-treated with DMSO, brazilin (10 µM) or dimethyl fumarate (DMF) (30 µM), a known inducer of NRF2 signalling [66], for 15 minutes before LPS priming (1 µg ml^−1^, 6 h). As expected, DMF alone induced NRF2 accumulation (Figure 4C) and this effect was enhanced by the addition of LPS (Figure 4C_i__-ii_). DMF pre-treatment also significantly reduced pro-IL-1β (Figure 4C_i__,iii_) and IL-6 levels (Figure 4D; Figure S1A raw data) in response to LPS priming compared to DMSO treated controls, but did not affect LPS-induced NLRP3 expression (Figure 4C_iv_) or TNFα release (Figure 4E; Figure S1B raw data). Treatment with brazilin alone did not induce significant NRF2 accumulation (Figure 4C_i_). However, as observed with DMF treatment, brazilin strongly enhanced LPS-induced NRF2 accumulation (Figure 4C_i__-ii_) whilst significantly inhibiting LPS-induced pro-IL-1β (Figure 4C_i__,iii_) and IL-6 production (Figure 4D; Figure S1A raw data), without affecting TNFα release (Figure 4E; Figure S1B raw data). Brazilin also significantly reduced NLRP3 expression in LPS-primed cells (Figure 4C_iv_), as we observed previously (Figure 4A_i_).

These results show that brazilin can inhibit LPS-induced priming of the NLRP3 inflammasome, which may be a consequence of brazilin enhancing NRF2 transcription factor signalling.

### Brazilin significantly reduces NLRP3 inflammasome activation in a human iPSC-microglial cell line

Dysregulated NLRP3 inflammasome activation has been implicated in several central nervous system (CNS) disorders, including stroke, Alzheimer’s disease and Parkinson’s disease [19]. To inform upon the potential clinical translation of brazilin for such conditions, we next sought to examine the effect of brazilin on inflammasome activity in human microglia; the predominant CNS-resident immune cells. We generated iPSC-derived microglial-like cells according to previously established methods [59, 60] (see also Supp Methods). Firstly, we verified that our iPSC-microglia expressed the key markers IBA1, TREM2 and P2Y12 using immunocytochemistry and fluorescence microscopy (Figure S2A_i-iii_). We also confirmed the microglia were functional and could respond to inflammatory stimuli by demonstrating their ability to phagocytose pHrodo-labelled *E.coli* bioparticles (Figure S2B) and their release of various cytokines (TNFα, IL-6, IL-10) and chemokines (CXCL10) following stimulation with LPS (Figure S2C). We also confirmed that our iPSC-microglia can mount an inflammasome response to LPS and nigericin stimulation by transfecting the cells with lentiviral human ASC-GFP and visualising ASC speck formation (Figure S2D).

To explore whether brazilin could influence the NLRP3 inflammasome response, human iPSC-microglia were primed with LPS (100 ng ml^−1^, 16 h) followed by 30-minute treatment with brazilin (10-100 µM), MCC950 (0.1 µM) or DMSO control, before NLRP3 inflammasome activation with nigericin (10 µM, 2 h). Brazilin (≥10 µM) treatment significantly reduced both the amount of IL-1β released (Figure 5A; Figure S3 raw data) and the number of ASC specks formed (Figure 5B-C) in iPSC-microglia, indicating a significant inhibition of NLRP3 inflammasome activation. Although brazilin appeared to inhibit IL-1β release in a dose-dependent manner, this was not the case with ASC speck formation, as brazilin concentrations >30 µM did not cause any further reductions in speck numbers (Figure 5B).

**Figure 5.**
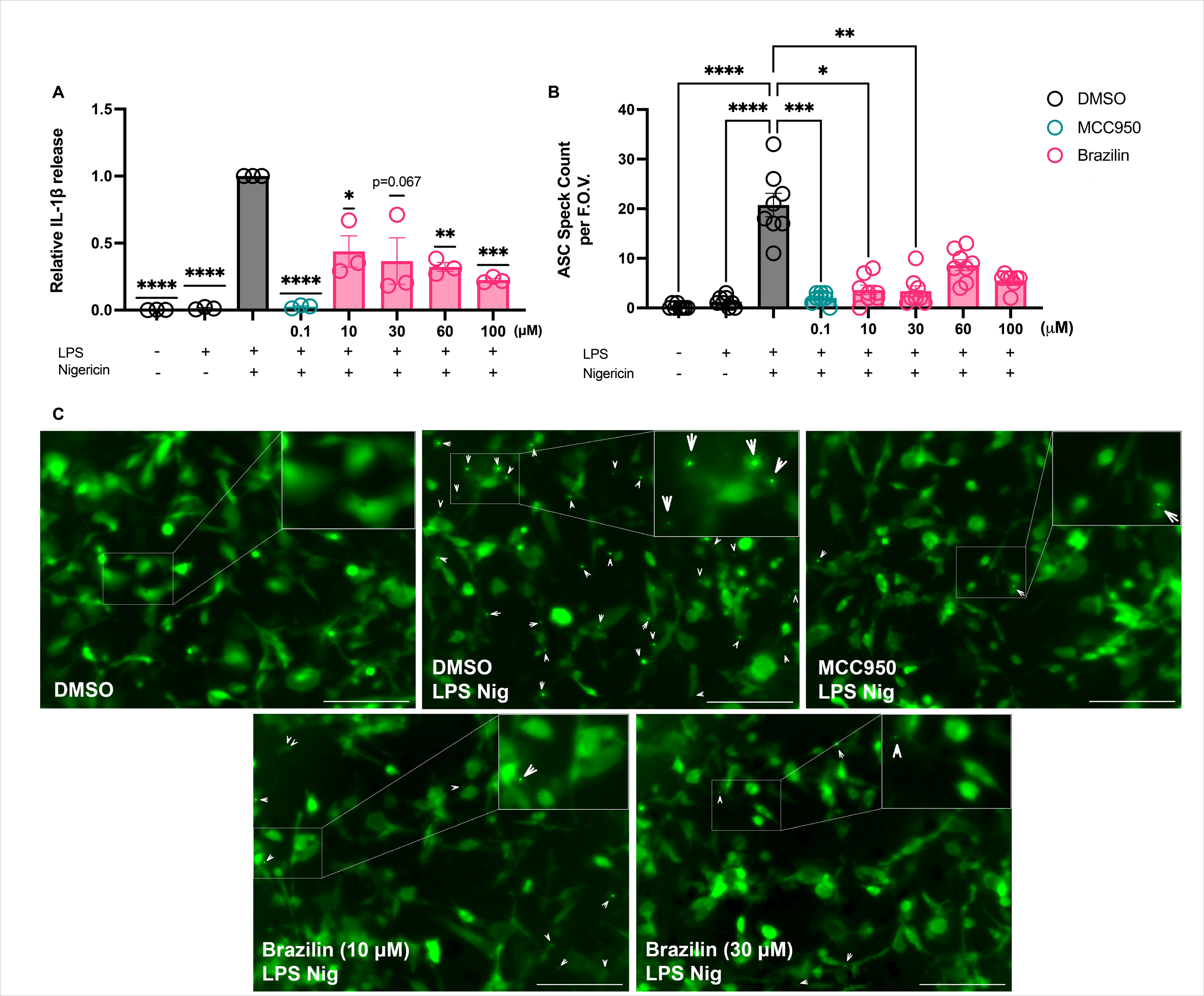
Brazilin significantly inhibits NLRP3 inflammasome activation by nigericin in human iPSC-microglia. Human iPSC-derived microglial cells expressing lentiviral hASC-GFP were primed with LPS (100 ng ml) or vehicle (16 h) before treatment with MCC950 (0.1 µM), brazilin (10, 30, 60 or 100 µM) or DMSO control (30 min). Nigericin was then added to activate the NLRP3 inflammasome (10 µM, 2 h). **(A)** Supernatant IL-1β was quantified using an IL-1β Human Luminex® Discovery Assay kit, expressed relative to DMSO control cells treated with LPS and ATP. See also Figure S3 for IL-1β release data without normalisation. N=3 biological repeats/group. **(B)** ASC specks were quantified via blinded manual speck counts from two images (field of view; F.O.V.) obtained from four biological replicates of each treatment condition (N=8). **(C)** Representative images used for ASC speck counting. ASC specks are indicated by arrow heads and a zoomed in region is placed in the top right corner of each image, indicated by a white box. Images were captured using SX5 incucyte imaging system (20X phase, green GFP fluorescence). Scale bars represent 200 µm. Statistical analyses following normality testing: (A) one-sample t-Tests, comparing means to a hypothetical mean of 1.0. (B) Kruskal-Wallis test with Dunn’s multiple comparisons. All data are expressed as mean ± SEM. *p<0.05, **p < 0.01, ***p < 0.001, ****p<0.0001.

### Brazilin significantly inhibits the NLRP3 inflammasome response to acute inflammation *in vivo*

To investigate whether brazilin treatment could inhibit the NLRP3 inflammasome response *in vivo*, mice were treated with a single i.p. dose of LPS, immediately followed by a single i.p. dose of brazilin (50 mg/kg), MCC950 (20 mg/kg) or vehicle (1% DMSO in PBS). After 2 hours, mice were stably anaesthetised and either PBS control or ATP was administered i.p. for 15 minutes under terminal anaesthesia, the latter used to activate the NLRP3 inflammasome. Peritoneal lavage was then performed and the fluid used for cytokine analyses (Figure 6A). As expected, vehicle pre-treated mice that received both LPS and ATP had elevated IL-1β levels in the peritoneum, compared to vehicle pre-treated mice that received LPS only (Figure 6B), indicating an activated NLRP3 inflammasome response. However, mice pre-treated with either brazilin or MCC950 had significantly lower levels of peritoneal IL-1β in response to LPS and ATP administration, compared to the vehicle-treated group (Figure 6B). This indicated an inhibition of the inflammasome response by both MCC950 and brazilin *in vivo*. Neither brazilin nor MCC950 significantly altered peritoneal levels of TNFα (Figure 6C) or IL-6 (Figure S4) evoked by LPS and ATP.

**Figure 6.**
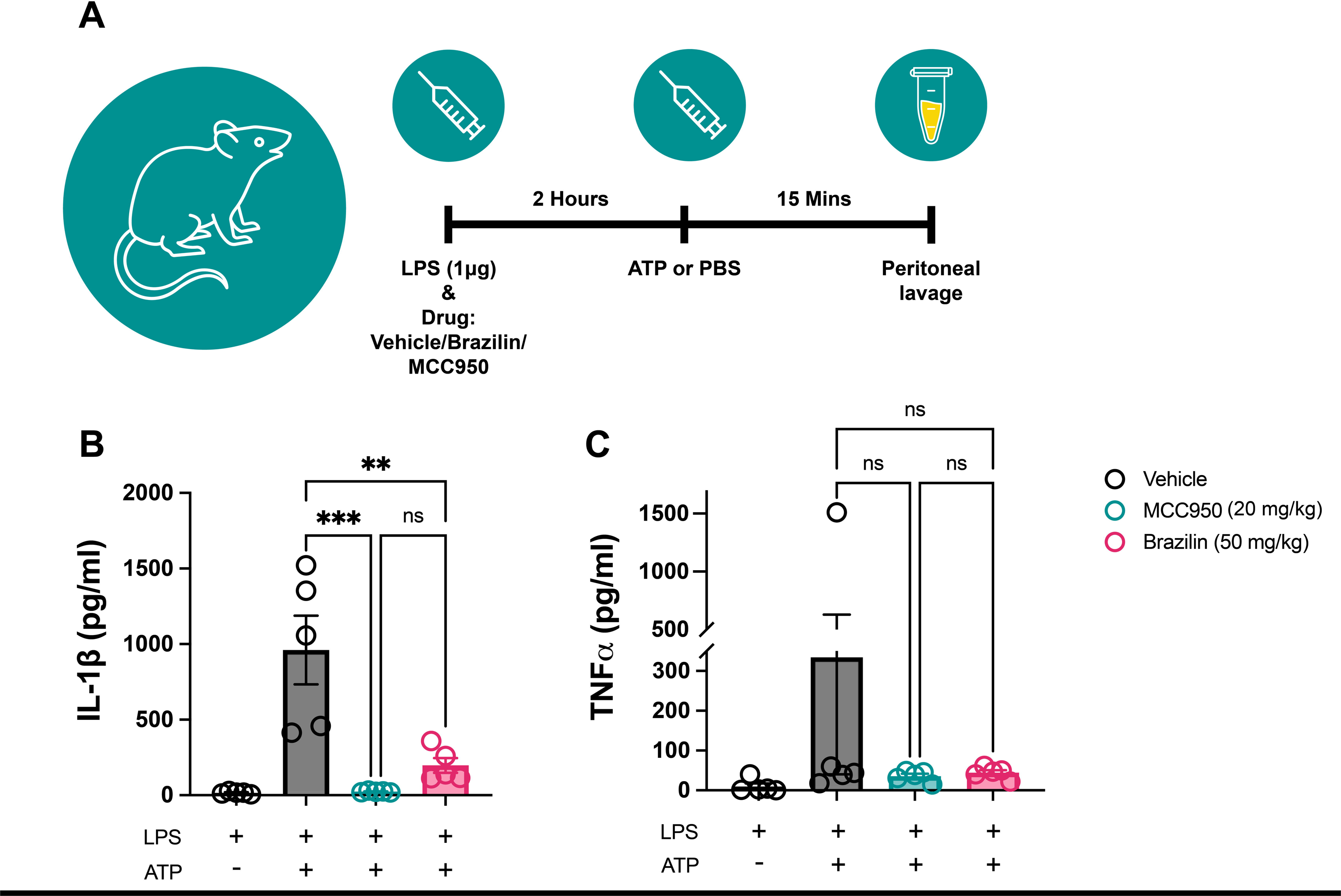
Pre-treatment with brazilin can significantly attenuate the NLRP3 inflammasome response in vivo. **(A)** Schematic of the treatment protocol and sample collection from male C57 mice. Intraperitoneal (i.p.) administration of either brazilin (50 mg/kg, 2h) or MCC950 (20 mg/kg, 2h), alongside LPS (i.p., 1 μg, 2h), **(B)** significantly attenuated subsequent IL-1β release in the peritoneum in response to ATP administration (i.p., 100 mM in PBS (500 µl/mouse), 15 min), compared to vehicle (1% DMSO in PBS) treated mice. Neither brazilin nor MCC950 treatments had any effect on peritoneal **(C)** TNFα release in response to LPS and ATP administration. See also Figure S4 for peritoneal IL-6 data. Data are presented as mean ± SEM. N=5 mice per treatment group. Statistical analyses following normality testing: (B) one-way ANOVA with Tukey’s post-hoc comparisons, (C) Kruskal-Wallis test with Dunn’s post hoc comparisons were used to assess the effect of drug treatment between groups treated with both LPS and ATP. *p<0.05, **p < 0.01 and ***p < 0.001. DMSO, dimethylsulfoxide; LPS, lipopolysaccharide; ATP, adenosine triphosphate; i.p., intraperitoneal.

## Discussion

The NLRP3 inflammasome is a cytosolic protein complex that drives pro-inflammatory cytokine production and is an executioner of lytic cell death. Inflammasome responses can serve as a rapid cellular defence mechanism to limit tissue damage and/or spread of infections, however, when dysregulated can become a facilitator or even instigator of disease progression. Molecular inhibition of the NLRP3 inflammasome has translational relevance for numerous conditions involving acute or chronic inflammation. However, no NLRP3 inflammasome inhibitors have yet been approved for therapeutic use in the clinic, with toxicity concerns prohibiting the progression of some compounds through clinical trials. As such, there remains a crucial need to continue developing new, tolerable inhibitors and there has been growing success in identifying lead compounds amongst natural products [39, 67, 68].

Here we have identified that the natural product brazilin can significantly inhibit the NLRP3 inflammasome response in cultured mouse macrophages and human iPSC-derived microglia. Our experimental data suggest that brazilin exerts a dual inhibitory effect upon the canonical NLRP3 inflammasome, whereby it can inhibit both the priming and activation steps of the pathway. By pre-treating murine BMDMs with brazilin, the expression of pro-IL-1β, IL-6 and NLRP3 was significantly reduced in response to priming with bacterial LPS. These findings are in line with previous studies that report brazilin to supress the pro-inflammatory response of cultured macrophages to LPS [31–33]. However, our data also indicate that brazilin inhibits LPS-induced expression of the NLRP3 protein.

Brazilin has been proposed to limit the cellular production of pro-inflammatory mediators through a general suppression of NF-κB signalling, by reducing NF-κB nuclear translocation [34] and/or the binding of NF-κB to DNA [31]. However, we found that brazilin treatment did not reduce the secretion of TNFα from macrophages, indicating that there was not global inhibition of NF-κB signalling in brazilin-treated cells. Elevated NRF2 signalling has previously been reported to underpin cardioprotective effects of brazilin against ischaemia-reperfusion injury [69]. NRF2 is a transcription factor which is known to inhibit the expression of NF-κB secondary response genes, including IL-1β and IL-6 [63], without inhibiting the expression of NF-κB primary response genes such as TNFα [66]. Additionally, brazilin has been shown to induce the expression of heme oxygenase-1 (HO-1) in cultured cells [32, 70] which may also be due to brazilin-mediated NRF2 activation, given that HO-1 is a target gene of NRF2 [64, 71]. Therefore, we investigated the accumulation of NRF2 in BMDMs following LPS stimulation and indeed found that this was enhanced by brazilin treatment. Therefore, we propose that in the context of inflammation, brazilin may inhibit the expression of specific LPS-induced gene subsets by increasing NRF2 signalling, which consequently contributes to an inhibition of NLRP3 inflammasome priming. However, the ability of brazilin to reduce NLRP3 expression after LPS priming may be independent of NRF2 signalling, as the NRF2 inducer DMF does not affect LPS- induced NLRP3 production [66].

Brazilin can also inhibit NLRP3 activation, via a mechanism that is reversible and independent of its influence on inflammasome priming. Treatment of LPS-primed BMDMs with brazilin was sufficient to inhibit processes downstream of NLRP3 activation, namely the maturation of caspase-1, gasdermin-D and IL-1β proteins, without altering the upstream production of pro-IL-1β. Brazilin also significantly reduced the formation of ASC specks and the release of IL-1β from human iPSC-microglia in response to nigericin. MCC950 and the analogue NP3-146 maintain the NLRP3 protein in an inactive conformational state by binding to a specific site within the NLRP3 NACHT domain [22–24]. Our computational modelling predicts that brazilin may also directly interact with this inhibitor binding site. Furthermore, our biological assays showed that brazilin was able to inhibit NLRP3 activation evoked by diverse signalling pathways, dependent or independent of potassium ion efflux, and brazilin had no inhibitory effect upon the activation of the AIM2 or NLRC4 inflammasome. Collectively, these findings support our hypothesis that brazilin may interact directly and selectively with the NLRP3 sensor protein to inhibit inflammasome activation. Through direct binding, brazilin may mediate degradation of the NLRP3 protein, which may explain the inhibition of LPS-induced NLRP3 expression and/or may contribute to the slight reduction in NLRP3 levels and inhibition of the inflammasome activation step that we report. However, further experiments are needed to address these hypotheses.

Brazilin has a lower potency to inhibit the NLRP3 inflammasome in BMDMs (IC_50_ 1.98 μM) compared to MCC950 (IC_50_ 7.5 nM) [20], which may be partially due to an oxidation of brazilin to form the red-pigmented brazilein product, which our computational modelling predicts to have a much lower binding affinity for NLRP3 than brazilin. The brazilin molecule could serve as a scaffold for the development of novel structural analogues with improved stability and potency to inhibit the NLRP3 inflammasome. Nevertheless, we have also shown that brazilin treatment can significantly attenuate peritoneal IL-1β release in response to LPS and ATP administration in mice, confirming that brazilin in its current form is sufficiently stable and active to inhibit an acute NLRP3 inflammasome response *in vivo*.

Brazilin is the most abundant constituent of heartwood from the *Haematoxylum brasiletto* and *Caesalpinia sappan* tree species. While there are no commercially licensed products containing purified brazilin, heartwood extracts from these trees have long been used as colouring agents in foods and beverages and as medicinal remedies [27, 28]. This would suggest that brazilin is likely safe for human consumption; however, thorough preclinical and clinical toxicology investigations are still required. While safety profiling was not an aim for our experiments, we can report that brazilin was not cytotoxic to cultured BMDMs (≤ 30 μM), which is supported by previous *in vitro* experiments [31–33]. Brazilin also had no obvious adverse effects on the appearance, behaviour or survival of the mice in our in vivo study, although their exposure to the treatment was short (2 hours) and no formal post-mortem analyses of organs were performed. We found that brazilin may enhance TNFα release from LPS-primed BMDMs, which may be a concern as TNFα is itself known to drive a range of inflammatory diseases [72]. However, brazilin treatment did not enhance peritoneal TNFα levels beyond that associated with the inflammation itself in our mouse model. It is also worth noting that brazilin had no inhibitory effects on the AIM2 inflammasome response to DNA or the NLRC4 inflammasome response to bacterial flagellin in cultured BMDMs, which may be an important attribute when considering such a compound as a potential immunomodulatory therapeutic. However, further experiments are required to confirm this in vivo.

In conclusion, we have identified that the natural product brazilin can exert a dual inhibitory effect over both the priming and activation steps of the canonical NLRP3 inflammasome pathway. Our results indicate that brazilin can enhance NRF2 signalling, which may contribute to the inhibition of inflammasome priming, while our computational modelling suggests that brazilin may also bind directly to the NLRP3 protein to inhibit inflammasome activation. We have confirmed that brazilin can significantly inhibit the NLRP3 inflammasome response in cultured murine macrophages, human microglial cells, and in a mouse model of acute inflammation. Our results encourage further evaluation of brazilin as a promising therapeutic agent for NLRP3-related inflammatory diseases. Moreover, this small-molecule may serve as a scaffold for the development of new selective NLRP3 inflammasome inhibitors.

## Significance

Inhibition of the NLRP3 inflammasome is a promising therapeutic strategy for targeting inflammation-associated pathology in a variety of human conditions. With no NLRP3 inflammasome inhibitors currently approved for clinical therapeutic use, there remains a great need to identify and develop new inhibitor compounds which have a low risk of adverse effects. In this manuscript, we provide compelling new evidence from in vitro and in vivo mouse experiments that the natural product brazilin is an NLRP3 inflammasome inhibitor. Brazilin is the major constituent of heartwood from several tree species and may have a favourable safety profile. Extracts from this heartwood have long been used as traditional medicines and we found no indications of brazilin-associated toxicity in our experiments, which is supported by previous publications. Importantly, we also show that brazilin can attenuate the NLRP3 inflammasome response in a human iPSC-microglial cell line, which implicates the translational relevance of our discoveries. We also describe the generation and utilisation of this human iPSC-microglial cell line to perform inflammasome assays in this manuscript, which we believe provides a powerful tool for the field of neuroimmunology research. Despite having a very different molecular structure to MCC950, our computational modelling predicts that brazilin may inhibit NLRP3 activation by binding to the same site of the protein. Therefore, this natural compound may offer a new scaffold for the design of derivative compounds with improved potency to inhibit the NLRP3 inflammasome whilst evading cellular toxicity.

## Acknowledgements

The authors thank Graham Coutts for assistance in the harvest of primary BMDMs for these experiments, Victor Tapia for training in cellular models and Robert White for assistance in figure design. This work was supported by the Medical Research Council (MRC) Doctoral Training Partnership [MR/N013751/1] PhD Programme at the University of Manchester. D. B. and C. H. were funded by an MRC grant [MR/T016515/1]. The authors would like to acknowledge the assistance given by Research IT and the use of the Computational Shared Facility at The University of Manchester.

## Author Contributions

E. M. and P. R. K. conceived the project and E. M. performed most of the experiments and analysed the data. All computational modelling was performed and transcribed by S. E., with assistance from R. A. B. and S. F. The iPSC line was obtained with informed consent and with the approval of Wellcome Sanger Institute. L. F. and E. J. characterised and performed all experiments using the iPSC cell line. C. H. assisted in the experimental design for testing the impact of brazilin treatment on NrF2 signalling. J. G. and C. B. L. assisted with intraperitoneal dosing of mice. E. M. and S. E. wrote the manuscript, and all the authors contributed to the editing.

## Supplemental videos

**Video S1 MD trajectories over 40ns MD simulation of S-brazilin-NACHT,** related to Figure 1. The video shows the interactions between S-brazilin (grey) and the inhibitor binding site of NACHT domain. Walker A (WA) site is coloured green.

