## Supplemental Materials for "Brazilin is a Natural Product Inhibitor of the NLRP3 Inflammasome"

**Supplemental Methods**

**Human iPSC-microglia characterisation**

For immunocytochemistry experiments, iPSC-microglial cells (11-14DIV) were washed with ice-cold PBS and fixed with 4% paraformaldehyde for 30 minutes. Following fixation, cells were permeabilized using Triton X-100 (0.2%)/PBS and immunolabeled with antibodies against IBA1 (FUJIFILM WAKO 019-19741, 1:300) and TREM2 (R&D Systems AF1828, 1:300) in 10% donkey serum/PBS. For P2Y12 labelling (Alomone APR-020, 1:100), cells were labelled following fixation without any permeabilization step. Primary antibodies were revealed with a 1 h incubation with Alexa Fluor secondary antibodies (1:1000, 10% donkey serum/PBS). Cells were incubated with DAPI (4’, 6-diamidino-2’-phenylindole, dihydrochloride) (Thermofisher, D1306) (1:1000, 1 mg/ml in water) during the penultimate wash. Confocal images were acquired using the Opera Phenix High-Content Screening System (PerkinElmer).

For the fluorescent ASC speck assay, cells were fixed in 4% PFA as described above, after which a Wheat Germ Agglutinin (WGA) cell mask (Invitrogen, W32466; 1:200 in PBS) and Hoescht nuclear stain (Invitrogen, 62249; 1:4000 in PBS) were added. Cells were then washed twice prior to confocal imaging.

For phagocytosis experiments, iPSC-microglia (11-14 DIV) were pre-treated with 3uM Cytochalasin D (Sigma) or DMSO control for 30 mins then incubated with 0.1 mg/ml pHrodo-labelled E.coli Bioparticles (Sartorius). Cells were imaged using a Incucyte cell imaging system every hour for 24 hours (10X, Phase and green fluorescence). Using IncuCyte software, image masks for the fluorescent signal were generated and quantified.

For cytokine release experiments, iPSC-microglial cells (11-14DIV) were stimulated with 100 ng/ml of LPS. Supernatants were harvested after 24 hours and analysed with a custom Human Luminex® Discovery Assay kit (R&D Systems) to examine secretion of TNF-⍺, IL-6, IL-10, and CXCL10 following stimulation.

**Supplemental Figures and Tables**

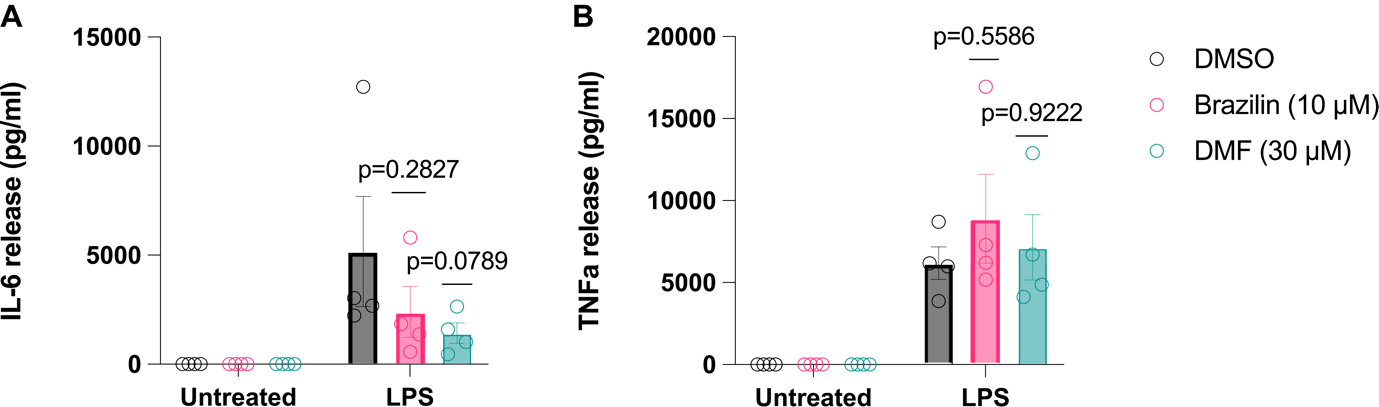

**Figure S1	Supernatant cytokine release from primary BMDMs after LPS priming**, related to Figure 3. BMDMs were treated with vehicle (DMSO), brazilin (10 µM) or DMF (30 µM) for 15 min. LPS (1 µg ml^−1^, 6 h) was then added to the wells to induce priming. Supernatants were analysed by ELISA for **(A)** IL-6 (N=4) and **(B)** TNFα (N=4) release. Data show absolute cytokine release (pg ml^-1^). Data are presented as mean ± SEM, each data point (‘N’) representing a biological repeat. Statistical analyses following normality testing: (A) Kruskal-Wallis test with Dunn’s post hoc comparisons, LPS+ data; (B) one-way ANOVA with Tukey’s post hoc comparisons, LPS+ data. *p<0.05. BMDMs, bone marrow-derived macrophages; DMSO, dimethylsulfoxide; DMF, dimethyl fumarate.

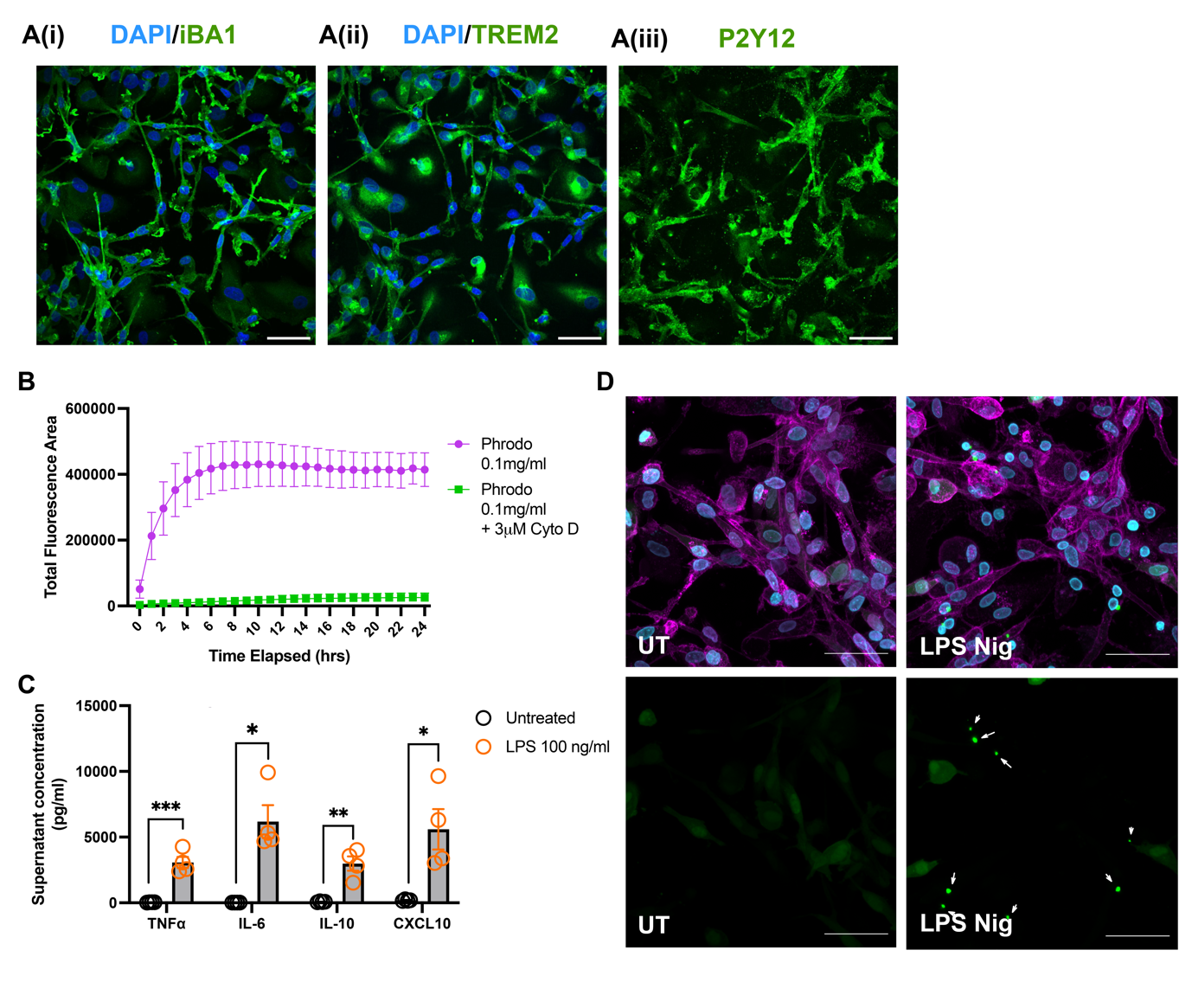

**Figure S2	Characterisation of human iPSC-derived microglia demonstrates they are functional and respond to inflammatory stimuli.**, related to STAR methods. (**A)** Representative fluorescent confocal microscopy images of iPSC-derived cells expressing markers typical of microglial cells: (Ai) iBA1, (Aii) TREM2 and (Aiii) P2Y12 (all green GFP). Nuclei stained with DAPI (blue). Scale bars represent 50 μM. **(B)** iPSC-microglia display the ability to phagocytose. Here shown is engulfment of pHrodo labelled E.coli bioparticles, which fluoresce once inside acidic phagosomes. Cytochalasin D (3μM) was added as a negative control to inhibit phagocytosis (n=4). **(C)** iPSC-microglia respond to LPS, releasing cytokines and chemokines. Measurement of supernatant cytokines/chemokines was performed using Luminex technology (n=4). **(D)** The formation of ASC specks can be seen in iPSC-derived cells transfected with lentiviral hASC-GFP. ASC specks are not present in untreated (UT) iPSC-microglia, but specks form in response to treatment with LPS (100 ng ml^-1^, 16 h) followed by nigericin (10 μM, 2 h). Arrow heads point to ASC specks. Wheat Germ Agglutinin (WGA) cell mask (magenta) and Hoescht nuclear stain (blue) are also shown. Scale bars represent 50 μM. All images were acquired using an Opera Phenix fluorescent confocal microscope.

Data are expressed as mean ± SEM. (C) Unpaired t-tests were used to compare TNFa, IL-10 and CXCL10 release between untreated and LPS treated cells. A Mann-Whitney test was used to compare IL-6 release between untreated and LPS treated cells. Each datum represents a biological repeat (‘n’).

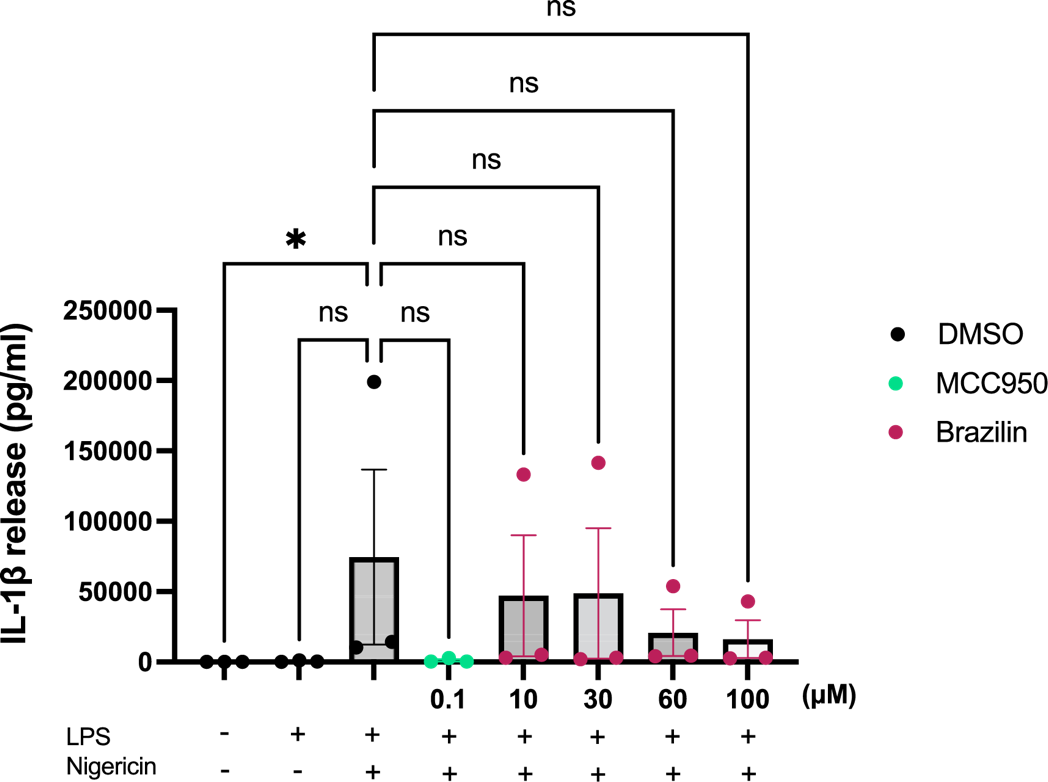

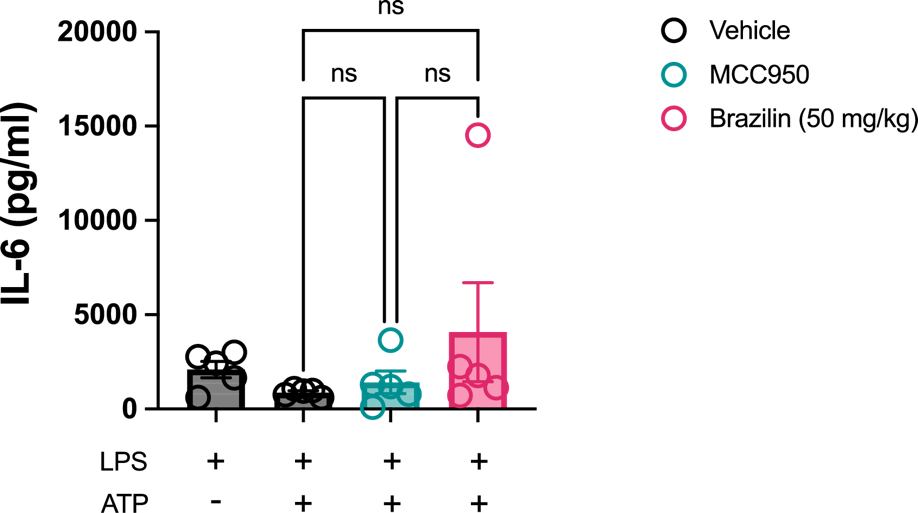
**Figure S3	Supernatant IL-1β released from human iPSC-microglia,** related to Figure 5. Cells were primed with LPS (100 ng ml^-1^) or vehicle (16 h) before treatment with MCC950 (0.1 µM), Brazilin (10, 30, 60 or 100 µM) or DMSO control (30 mins). Nigericin was then added to activate the NLRP3 inflammasome (10 µM, 2 h). Supernatant IL-1β was quantified using an IL-1β Human Luminex® Discovery Assay kit. Kruskal-Wallis test with Dunn’s multiple comparisons. All data are expressed as mean ± SEM. *p<0.05. Each data point (‘N’) represents a biological repeat.

**Figure S4	Pre-treatment with brazilin does not alter peritoneal IL-6 release in response to LPS and ATP administration,** related to Figure 6.

Male C57 mice (N=5 mice/group) received a single i.p. dose of either vehicle (1% v/v DMSO in PBS), brazilin (50 mg/kg) or MCC950 (20 mg/kg), alongside LPS (1 μg; i.p.). After 4 hours, mice received a single i.p. dose of ATP (100mM in PBS) or vehicle (PBS), 15 minutes prior to peritoneal lavage.

Data are presented as mean ± SEM. One-way ANOVA with Tukey’s post-hoc comparisons, to assess the effect of drug treatment between groups treated with both LPS and ATP. DMSO, dimethylsulfoxide; LPS, lipopolysaccharide; ATP, adenosine triphosphate; I.P., intraperitoneal.

**Table S1.** Average interatomic distances (in Å) between NACHT residues and selected carbon atoms of S-brazilin and S-brazilein from the 40 ns MD of ligand-NACHT complexes. The initial distance in docked poses was measured using MOE. Standard deviations in parentheses.

| 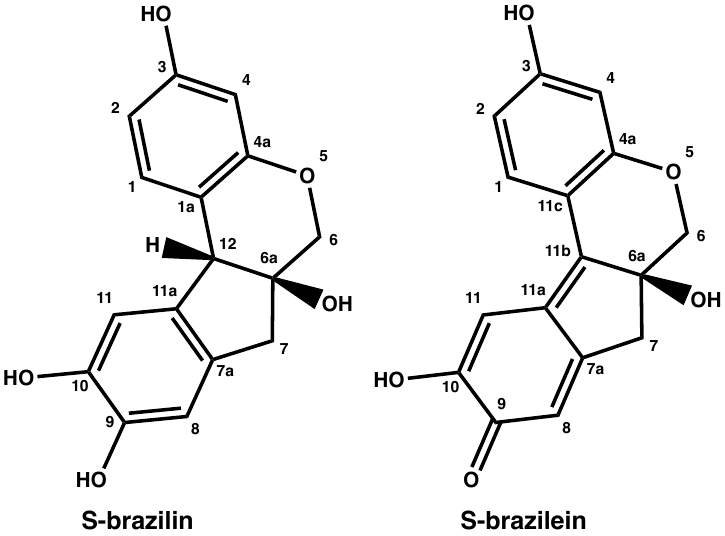 | | | | |
| --- | --- | --- | --- | --- |
| Distance | S-brazilin | | S-brazilein | |
|  | Initial | MD | Initial | MD |
| C2-Leu413C_β_ | 3.85 | 4.08 (0.27) | 3.60 | 5.04 (0.92) |
| C3-Leu413C_δ1_ | 4.43 | 4.40 (0.44) | 3.89 | 4.84 (1.00) |
| C4a-Ile234C_δ1_ | 3.89 | 4.16 (0.43) | 4.71 | 5.59 (1.13) |
| C7-Ile151C_δ1_ | 5.28 | 4.26 (0.36) | 4.59 | 4.76 (0.70) |
| C8-Tyr381C_ε2_ | 4.27 | 4.36 (0.36) | 3.99 | 4.38 (0.41) |
| C9-Tyr168C_δ1_ | 4.03 | 4.69 (0.32) | 4.10 | 4.42 (0.39) |
| C10-Ile234C_γ2_ | 4.12 | 3.88 (0.29) | 3.73 | 4.46 (0.44) |
| C11-Ile234C_γ1_ | 4.39 | 3.87 (0.30) | 3.92 | 4.82 (0.59) |
| C11-Pro412C_β_ | 3.09 | 3.58 (0.24) | 3.64 | 4.22 (0.57) |
